# Mitigating Local Over-fitting During Single Particle Reconstruction with SIDESPLITTER

**DOI:** 10.1101/2019.12.12.874081

**Authors:** Kailash Ramlaul, Colin M. Palmer, Christopher H. S. Aylett

**Affiliations:** Section for Structural and Synthetic Biology, Department of Infectious Disease, Faculty of Medicine, Imperial College Road, South Kensington, London, SW7 2BB, United Kingdom; Scientific Computing Department, Science and Technology Facilities Council, Research Complex at Harwell, Didcot, OX11 0FA, United Kingdom

**Author notes:** These authors contributed equally to this work (Kailash Ramlaul, Colin M. Palmer).

**Keywords:** Cryo-EM, Local resolution, Noise suppression, Real-space filter, Over-fitting

## Abstract

Single particle analysis of cryo-EM images enables macromolecular structure determination at resolutions approaching the atomic scale. Experimental images are extremely noisy, however, and during iterative refinement it is possible to stably incorporate noise into the reconstructed density. Such “over-fitting” can lead to misinterpretation of the structure, and thereby flawed biological results. Several strategies are routinely used to prevent the spurious incorporation of noise within reconstructed volumes, the most common being independent refinement of two sides of a split dataset.

In this study, we show that over-fitting remains an issue within regions of low local signal-to-noise in reconstructed volumes refined using the half-set strategy. We propose a modified filtering process during refinement through the application of a local signal-to-noise filter, SIDESPLITTER, which we show to be capable of reducing over-fitting in both idealised and experimental settings, while maintaining independence between the two sides of a split refinement. SIDESPLITTER can also improve the final resolution in refinements of structures prone to severe over-fitting, such as membrane proteins in detergent micelles.

## 1. Introduction

### 1.1 Improved versions of the iterative projection matching approach underlie most current single particle 3D reconstruction techniques

Technological developments are enabling single-particle reconstruction of macromolecules to increasingly high resolutions (Frank, 2017; Elmlund *et al*, 2017), providing a viable alternative to crystallography for larger (>100 kDa) complexes. The success of electron cryo-microscopy (cryo-EM) as a structure determination technique is underpinned by the process of single-particle analysis, which entails the three-dimensional (3D) reconstruction of macromolecular electron density from thousands or millions of 2D projection images of single particles (Elmlund & Elmlund, 2015; Carazo *et al*, 2015; Vilas *et al*, 2018b; Lyumkis, 2019).

Reconstruction of the macromolecular structure of interest is usually carried out in reciprocal space and relies on the Fourier projection (or central slice) theorem, which states that the 2D Fourier transform of an object’s projection is equivalent to a slice through the centre of the Fourier transform of the projected object in 3D (Bracewell, 1956). The correct alignment of each particle is essential for the reconstruction from 2D to 3D space, and thus the accurate estimation of the angular and positional parameters represents the defining problem of 3D reconstruction.

Most current computational procedures used to achieve alignment are derived from improvements to the projection matching process (Penczek *et al*, 1994), in which experimental projections are compared to *in silico* projections of an initial 3D reference map, and assigned the orientation parameters of the *in silico* projection based on their calculated similarity. Direct assignment, maximum likelihood, Bayesian *maximum a posteriori*, and other less well-defined statistical approaches have been applied to improve the performance and reduce the bias inherent to this process (Scheres, 2012a; Carazo *et al*, 2015). Iterated reconstruction and angular assignment allows the optimisation of the parameters assigned to each projection, leading to a stable and representative 3D reconstruction, providing that the sample is sufficiently homogenous and that the initial 3D reference was sufficiently accurate to allow convergence.

### 1.2 The independent 3D refinement of two halves of a split dataset is the most common method currently used to avoid over-fitting

Cryo-EM data are exceptionally noisy, an issue to which several factors contribute. Firstly, and chiefly, it is necessary to limit the electron dose used to acquire each image, because of the extensive radiation damage caused, resulting in “shot noise” due to stochastic sampling of the electron scattering probability distribution. Secondly, conformational and compositional variations between particles (including those due to accumulated radiation damage) result in heterogeneity. Finally, a number of other errors, such as incorrect parameter estimation (for example in determination of the CTF, or contrast transfer function), optical aberrations, and temporal variations, (e.g. uncorrected beam-induced motion), all contribute to further degradation of the signal, resulting in data with an extremely low signal-to-noise ratio (SNR) (Liao & Frank, 2010; Penczek, 2010; Vilas *et al*, 2018b).

During iterative independent refinement both the noise and signal from the data will be incorporated into each successive pair of structures. It is essential to suppress noise before the next alignment step, since otherwise images may be aligned to the noise, incorporating it stably into successive structures and forming new features that are indistinguishable from signal. This phenomenon is termed over-fitting (Grigorieff, 2000; Scheres & Chen, 2012). Amplified noise may be perceived as a real feature, and therefore over-interpreted (Scheres, 2012, Chen *et al*, 2013). Because the SNR decreases with increasing resolution, this has historically been handled by global filtering of the output structures at a chosen resolution beyond which the structure is considered “too noisy”.

The widely-accepted procedure for resolution assessment during EM reconstruction is the calculation of the Fourier Shell Correlation (FSC) between two halves of a single dataset (Harauz & Van Heel, 1986; Rosenthal & Henderson, 2003; Scheres & Chen, 2012). The collected particle images are randomly split into two half sets, and each half refined separately, but using identical procedures, as one side of an independent pair of reconstructions (Fig. 1A) (Grigorieff, 2000; Scheres & Chen, 2012; Henderson *et al*, 2012). The cross-correlation between Fourier components in successive resolution shells of each half-map is then calculated (Fig. 1B). At low resolution, the correlation between half-maps is expected to be high (approaching 1); at high resolution, the correlation should oscillate around zero.

**Figure 1:**
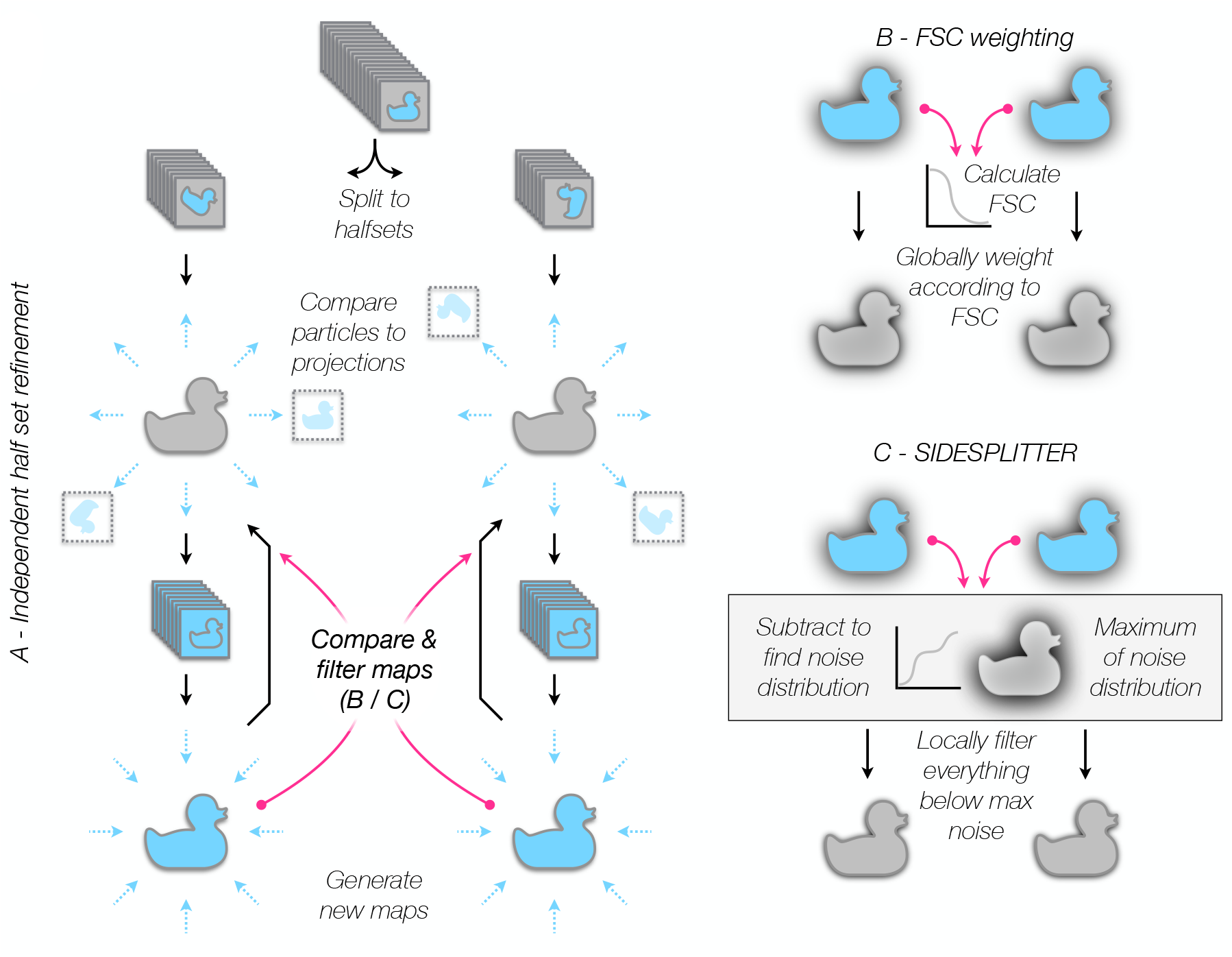
The application of SIDESPLITTER during the refinement process. Flow diagrams illustrating (A) pseudo-independent half-set projection matching refinement, (B) the application of FSC weighting as currently common in refinement, and the application of (C) SIDESPLITTER during such a refinement. Sources of information transfer between the pseudo-independent sides of the refinement are indicated with magenta arrows.

The resolution cut-off at which the reconstruction should be low-pass filtered remains a highly debated topic in the field (Van Heel & Schatz, 2005), however the most commonly used choice is an FSC value of 0.143 (Rosenthal & Henderson, 2003). Ideally, the measure of agreement needed for cryo-EM reconstructions is that between the experimental reconstruction, comprising both signal and noise, and the true structure, comprising only signal. Although a noiseless structure cannot be obtained, a theoretical estimate of this correlation can be calculated, which is denoted C_ref_. This is argued to represent the cryo-EM equivalent of the crystallographic figure of merit, and therefore a C_ref_ value of 0.5 (which occurs when the FSC value is 0.143) represents a resolution criterion consistent with crystallography (Rosenthal & Henderson, 2003).

Because the sides of the refinement are kept independent, in theory the FSC calculation between the two sides will be statistically correct, and filtering at the threshold resolution at each iteration should prevent noise incorporation at frequencies beyond the resolution cut-off.

### 1.3 Globally filtered reconstruction methods must be expected to over-fit local noise, as they cannot take account of local variations in SNR

Low-pass filtering based on a certain resolution cut-off will only exclude noise of a higher resolution from being incorporated into a refinement. The phenomenon of over-fitting has therefore become intimately linked to the evaluation of the resolution of cryo-EM reconstructions, where the resolution represents the maximum spatial frequency at which the information in the map is considered reliably interpretable as signal (Penczek, 2010). Whereas in crystallography, crystal packing of ordered units results in a relatively constant resolution throughout the entire electron density distribution, there is variability in cryo-EM reconstructions arising from heterogeneity in the particles used for reconstruction, non-uniform 3D reconstitution of Fourier components in Fourier space (Grigorieff, 2000) and inaccurate estimation of particle orientations (which causes a progressive degradation of resolution towards the edges of the reconstruction). The result is observed in real space as locally variable SNR within the reconstruction (Cardone *et al*, 2013). This effect is particularly pronounced in regions of a structure which display conformational flexibility or partial occupancy, and typically these regions suffer from lower interpretability as a result.

It has been shown previously that over-fitting occurs preferentially in regions of low SNR, i.e. where signal becomes indistinguishable from noise (Stewart & Grigorieff, 2004). Several methods for evaluation and treatment of local resolution in cryo-EM reconstruction have previously been proposed (Stewart & Grigorieff, 2004; Cardone *et al*, 2013; Chen *et al*, 2013; Kucukelbir *et al*, 2014; Vilas *et al*, 2018a; Ramírez-Aportela *et al*, 2018), however none provide a way to minimise over-fitting during the reconstruction process.

### 1.4 Local SNR filtering can mitigate local over-fitting throughout the structure during the refinement process

The challenge in 3D reconstruction is to make the best use of the available signal without incorporating noise. Therefore, we should aim to maximise the contribution of the available signal at all spatial frequencies during refinement, without over-fitting in regions of lower SNR (and hence lower local resolution). Recently, we introduced a new de-noising algorithm, LAFTER, which reduces the contribution of noise to cryo-EM reconstructions using two sequential real-space filtering steps (Ramlaul *et al*, 2019). LAFTER appeared to us to be particularly promising as a filter to prevent over-fitting, since it reduces noise by an optimal amount as judged by comparison to C_ref_, which is the most common standard currently used during independent half-set refinement (Rosenthal & Henderson, 2003). LAFTER was not applicable to the over-fitting problem during 3D refinement however, as it shares information between the two half-set reconstructions and therefore violates the requirement for the two halves to be independent.

In this paper, we present SIDESPLITTER, a heavily modified adaptation of the LAFTER SNR filter optimised to process both *sides* of a *split* refinement, which we successfully integrate into a 3D refinement workflow. SIDESPLITTER maintains the independence between the signal in the two sides of the refinement, sharing only the statistical properties of the noise distribution (Fig. 1C). We show that over-fitting is more pronounced in regions of lower local SNR, using both experimental reconstructions and synthetic datasets with explicitly-defined local resolution gradients. We further show that the application of the SIDESPLITTER noise-minimisation algorithm during iterative 3D refinement minimises over-fitting in poorly-resolved regions whilst retaining strong interpretable signal, and improving resolution in structures with severe over-fitting problems.

## 2. Methods

### 2.1 Justification and aims

Our first key aim is to minimise the residual noise during the refinement process, which biases the alignment on both sides of a split refinement and thereby results in over-fitting. We aim in particular to reduce residual noise within regions of lower local SNR that are not currently protected by the global filtering approaches in widespread use. Our second key aim is to maintain the independence between the two sides of the split refinement, since violation of this independence would lead to overestimation of the resolution of the reconstruction and would risk global over-fitting.

The first aim requires a local filter that is capable of suppressing noise to the greatest possible extent. The second requires that we avoid the use of a shared local window or a shared resolution map for both half-sets, as these would readily generate artefactual correlations between the sides of a split refinement (Supp. Fig. 1). Only global noise information can be shared without generating spurious correlations, and therefore a filter must either be capable of estimating SNR from the map alone (very difficult in masked refinements as no regions of pure noise are available), or must use only global statistics about the similarities and differences between the two sides for this purpose.

To achieve these two aims we have adapted our previous SNR filter based on local agreement (LAFTER) (Ramlaul *et al*, 2019), which has notable benefits for this application in that it suppresses noise to a greater degree than other local filters. To do this we have modified it to share only global statistics on the noise distribution in each shell, and no local statistics, between the independent sides. Therefore SIDESPLITTER will not result in any greater sharing of information between sides of the refinement than is the case for FSC weighting, which is already in regular use (Supp. Fig. 1) (Grigorieff, 2000; Scheres & Chen, 2012; Henderson *et al*, 2012).

### 2.2 Necessary assumptions

We assume that the noise is statistically independent between the two half-sets of the refinement process, while any agreement between the sides represents signal. This assumption is necessary as the properties of the noise distribution are estimated from the difference between sides. It should be noted that this same assumption underlies current global filtering approaches during refinement.

Secondly, the signal is assumed to vary smoothly in real space. This is necessary as we operate upon isolated frequency bands from the reconstructed maps in Fourier space, and any discontinuities in the signal would affect all frequency bands. This assumption is justifiable as macromolecular structures are known to be smooth at all resolutions accessible to cryo-EM.

Finally, as we use the maxima of the overall noise distribution between the half-sets to define the extent of the noise within the refinement, the noise must be reasonably evenly distributed over the density. Regions of exceptionally strong noise would be expected to result in over-aggressive filtering, although this will not necessarily be detrimental to the refinement process. We note that aggressive filtering is emphatically a lesser evil than the alternative.

### 2.3 The SIDESPLITTER algorithm

SIDESPLITTER was derived from the LAFTER SNR-based local filter. Extensive modifications have been made to the original algorithm in order to maintain as much independence between the two sides of the refinement as possible. In order to make SIDESPLITTER compatible with modern refinement algorithms (Scheres, 2012b; Punjani *et al*, 2017), the power spectrum is normalised to that of the input volume but tapered according to the estimated SNR to provide a representative spectrum. The overall approach follows a similar, two-pass, pattern to LAFTER, first normalising resolution shells in order to allow the SNR to be evaluated independently of resolution, and then truncating the frequencies contributing to each voxel at the resolution at which the signal falls below the maximum of the noise.

Initially, the spectral power of each input half map is calculated by taking the Fourier transform of each map, and then summing the power in each resolution shell:

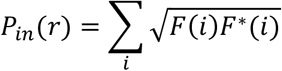

where *P*_*in*_(*r*) represents the power of the input map within the shell at resolution *r*, while *F*(*i*) and *F*\*(*i*) refer to the value and complex conjugate of a point *i* within the shell. The power spectrum for each half is stored, to be modified according to the SNR and reapplied at the end of the process so that the grey-scale can be maintained in the output half maps.

SIDESPLITTER then normalises the half-volumes. First, resolution shells are isolated from the two half volumes by band-pass filtering. The half volumes are transformed into Fourier space, and for each resolution shell, the Fourier coefficients are weighted using an eighth-order Butterworth band-pass filter (Butterworth, 1930):

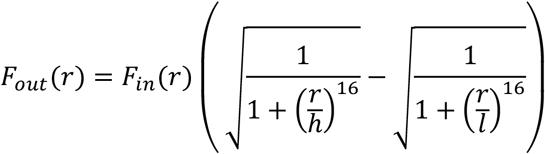

*F*_*in*_(*r*) and *F*_*out*_(*r*) represent the complex Fourier coefficients in the original transform and the band-passed output respectively, at radius *r*, while *h* and *l* represent the high and low cut-off frequencies. For each resolution shell, the two half volumes are then transformed to real space after the band-pass filter has been applied. The power of the combined map at this resolution, *T*, and the power of the noise, *N*, are calculated from the sums and differences of the voxel values respectively:

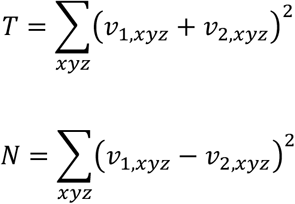

*v*_1,*xyz*_ and *v*_2,*xyz*_ represent the magnitude of the voxels from the two half-volumes at position *xyz*, and the sum is over all voxel positions within the mask (as provided by the user, or a simple spherical mask otherwise).

The proportional contributions of the noise and the signal to the total power are calculated as follows:

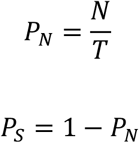

The power spectra (as calculated earlier) will be multiplied by *P*_*S*_ to weight them appropriately according to the SNR in each shell.

The voxel values in real space are normalised (to make them comparable for the second filter) by the resolution shell width and the root mean squared value of the total power at that resolution:

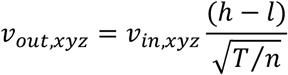

Incorporation of further high-resolution shells is terminated once the FSC within the mask falls below 0.143. After all resolution shells have been processed, the series of band-passed, noise-weighted maps for each half volume is summed in real space, combining the isolated resolutions to yield a pair of normalised half volumes, which should remain statistically independent over resolution, having only been scaled by simple multiplication in each shell.

In the filtering step the noise-suppressed half volumes from the first filter are transformed into the Fourier domain, and then each is low-pass filtered at every resolution that was considered in the previous step. Low-pass filtering is performed similarly to the band-pass filtering described above, using an eighth-order Butterworth response (Butterworth, 1930):

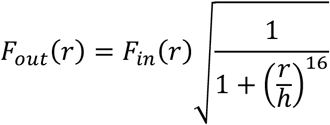

Each pair of low-pass-filtered half-maps is transformed back into real space. The observed maximum noise between half volumes is found as the greatest difference between corresponding voxels in the half volumes, for all voxel coordinates *xyz* within the masked region:

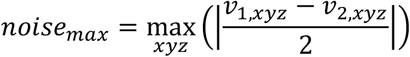

An expected upper bound on the maximum of the noise distribution, assuming the noise is normally distributed, is also calculated, according to the following formula, derived using Jensen’s inequality (Jensen, 1906) as is standard:

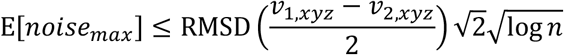

where RMSD 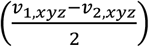 represents the root-mean-square deviation of the halved voxel differences within the mask, and *n* is the number of voxels considered.

Whichever noise bound is more conservative is used; the expected bound is a better estimate where the noise is close to being normally distributed, whereas the observed bound acts as a fall-back for cases in which the noise is strongly non-normally distributed, which is common when symmetry averaging operations have been applied. The noise values are halved in each case to account for the fact that they will be compared to voxel values in each half map separately.

Starting at the highest resolution considered, each voxel in each half volume is tested. If its value is greater than the noise bound at the current resolution, then that value is assigned to the corresponding voxel in that output half volume. If its value is lower than the noise maximum, the corresponding voxel in the output half volume is left un-assigned, and will be re-considered at the next (lower) resolution. Voxels that have already been assigned at higher resolution are excluded from consideration at lower resolutions, so the overall effect is that each voxel in each output half map is assigned to its value at the highest resolution at which its signal is greater than the maximum noise.

In order to preserve the grey-scale from the input, the output half maps are Fourier transformed and their power spectra calculated as before:

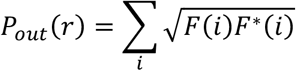

The output Fourier coefficients are then renormalized to the original power spectra and weighted by the SNR:

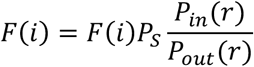

Finally, the output maps are transformed back to real space and passed into the 3D refinement program as reference volumes for it to use in its next iteration.

### 2.4 SIDESPLITTER reference implementation details

We provide a reference implementation of SIDESPLITTER as an optimised C99 program using FFTW3 for Fourier transformation (Frigo and Johnson, 2005) to maximise speed and portability. SIDESPLITTER operates upon MRC mode 2 format maps (Cheng et al., 2015), i.e. C float or FORTRAN real. Source code for the SIDESPLITTER reference implementation is available from the Imperial College Section for Structural Biology GitHub (github.com/StructuralBiology-ICLMedicine) under the GPL open source licence. SIDESPLITTER can be compiled for any POSIX-compatible operating system, and will also be made available in pre-compiled binary format for both Linux and Mac OS X as part of the CCP-EM suite (Burnley et al., 2017).

### 2.5 Generation of synthetic data for testing

For the synthetic test macromolecular structure, density for the AAA+ ATPase p97 was generated from an atomic model (PDB ID 1R7R; Huyton *et al*, 2003), which was used to create a benchmark synthetic dataset. Density from the molecular model was generated using `phenix.fmodel` from the PHENIX suite of programs (Adams *et al*, 2010) followed by the CCP4 suite program `fft` (Collaborative Computational Project, 1994). Each protomer of the model map was then masked individually and explicitly low-pass filtered to a given resolution stepwise around the ring (0.0125, 0.025, 0.05, 0.1, 0.2 and 0.35 cycles per voxel), creating a defined local resolution gradient around the ring. The SNR with resolution was explicitly maintained throughout using Gaussian noise. These locally filtered volumes were then modulated with a tau-factor falloff taken from the experimental SWR1-nucleosome dataset (Willhoft *et al*, 2018). The RELION utility `relion_project` was used to generate projections from the synthetic volumes, where the orientational distribution, CTF and noise parameters were taken from the experimental SWR1-nucleosome dataset (Willhoft *et al*, 2018) following a methodology similar to that previously reported for analysis of γ-secretase (Bai *et al*, 2015). The final projection images therefore represent a noisy, CTF-convoluted experimental dataset with a non-uniform distribution of projections exactly equivalent to the donor dataset.

### 2.6 Experimental datasets used for testing

Experimental datasets corresponding to EMD-9849 / EMPIAR-10264 (Lee *et al*, 2019), and EMD-4038 (Wilkinson *et al*, 2016) were kindly made available by T Nakane, Y Lee and DB Wigley, to test the applicability of the SIDESPLITTER refinement process within experimental refinement workflows.

### 2.7 3D reconstruction pipeline during SIDESPLITTER testing

Reconstructions were performed using RELION 3.0 (synthetic data) and an alpha version of RELION 3.1 (experimental data). All data were treated to an “auto-refinement” of half-sets in RELION, starting from known angular positions, but at low (50 Å) resolution, and otherwise with default parameters apart from the “`--solvent_correct_fsc`” flag, which was applied throughout, and a mask generated in RELION 3.0, which was applied with the “`--solvent_mask`” flag.

In the case of synthetic data, for which the angles needed examination on each cycle to maintain the correct subunit positions, RELION 3.0 was run for single iterations at a time, each called with the “`--continue`” flag. Between each iteration, SIDESPLITTER was applied to the unfiltered half maps output by the refinement job, and the “`_data.star`” files processed with a python script (provided alongside the SIDESPLITTER source code) that ensured that the particles had not moved to an adjacent subunit in the ring, by rotating particles that have deviated further than 30° from their known angle an additional 300° in the same direction.

For experimental data, through the use of an alpha version of RELION 3.1, we were able to make use of an additional new feature built into the `relion_refine` program. When called with an additional argument (`--external_reconstruct`), `relion_refine` calls an external program to perform reconstruction of the half maps after each iteration of 3D refinement. We used this as a hook to allow us to filter the half maps after they have been reconstructed and before the next iteration begins. A script that can be used to run SIDESPLITTER in the context of a RELION 3.1 refinement job is provided alongside the SIDESPLITTER source code.

## 3. Results

### 3.1 Synthetic data demonstrate that regions of low local resolution remain prone to residual over-fitting during single particle refinement of independent half-sets

Over-fitting remains an issue within regions that have a lower local resolution than those at the highest resolution in the reconstruction, even during independent half-set refinement. To demonstrate this, we generated synthetic data so that we could explicitly define and control the SNR of the underlying structure (Fig. 2). We generated map density from a molecular model with six-fold symmetry, and truncated the resolution of each subunit to a different resolution between 0.0125 and 0.35 cycles per voxel (Methods section 2.5). Knowing the exact properties of the underlying structure allows us to conclude that any correlation between the datasets beyond the expected resolution is due to noise retained during the refinement process, as there is known to be no initial signal to recover. These data were refined as normal in two independent half-sets, the only caveat being that the angular orientation was restricted to the 60° segment in which the projection was known to lie, to prevent the contribution of higher SNR information to the wrong segment.

**Figure 2:**
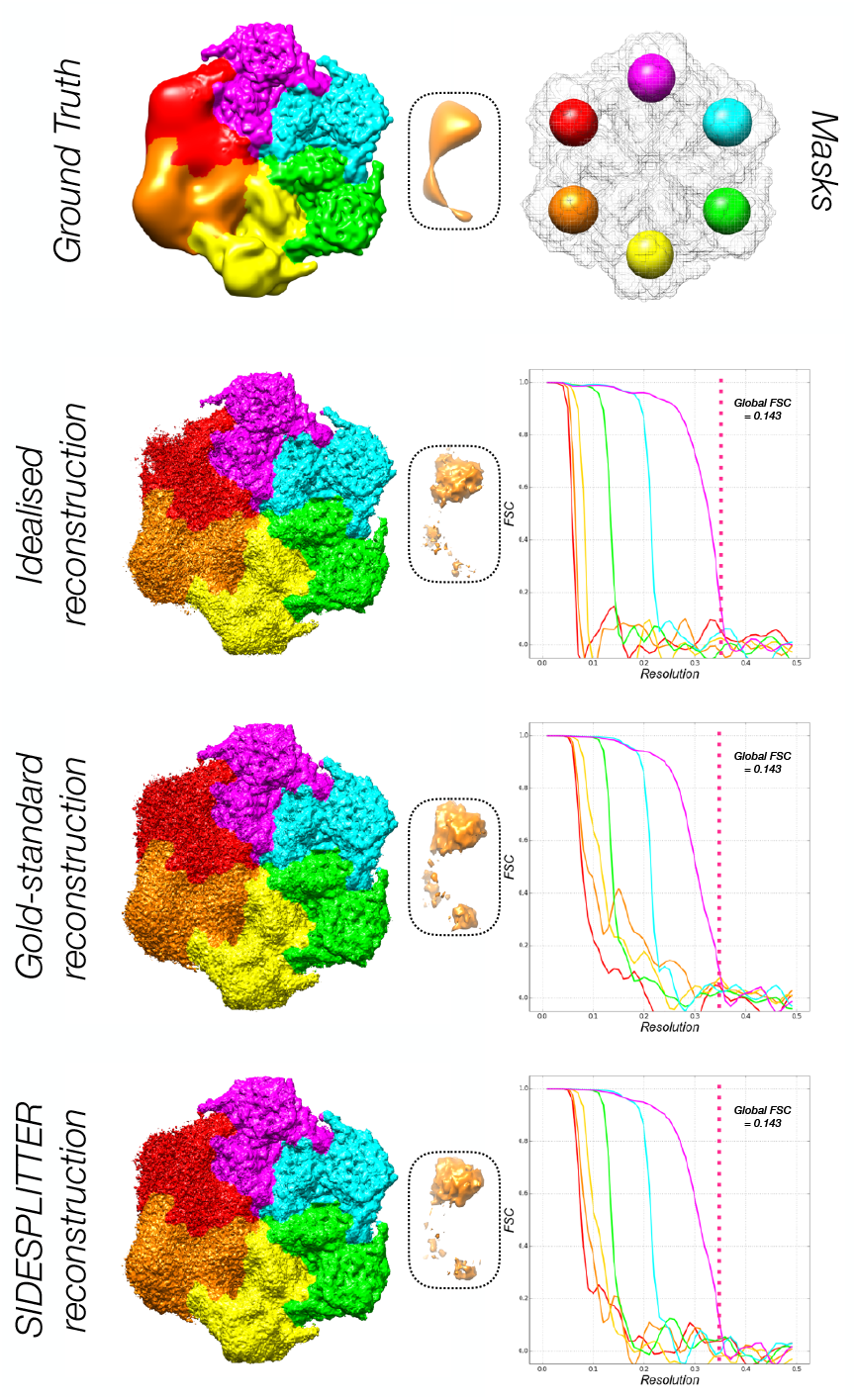
Refinement of synthetic data of known local resolution demonstrates that over-fitting occurs during independent refinement, and that SIDESPLITTER refinement reduces over-fitting. Panels indicate the ground-truth input, the output of idealised refinement against ground-truth, the output of standard independent refinement, and the output of refinement with SIDESPLITTER. Panels inset show density peaks in low local resolution. Over-refinement manifests as over-emphasis upon these peak regions. All volumes are shown as surfaces. The FSC curves between half-sets within soft spherical masks isolating part of each segment are inset, coloured according to the rainbow from red to purple (0.0125 to 0.35 voxels per cycle respectively). The corresponding masks are shown above, within the overall mask used during refinement (in grey).

While we observed no over-fitting beyond the known global resolution cut-off (0.35 cycles per voxel), there was evidence of visible features at higher resolution than the known resolution of the signal within the segments of low local resolution (Fig. 2, gold-standard reconstruction). These observations were confirmed by the FSC curves between masked regions in each segment. The FSC curves for the regions of low local resolution fall off from the known point of truncation, but exhibit residual correlation to higher resolutions (Fig. 2). For example, the orange curve remains significantly above zero out to almost 0.3 cycles per voxel, much higher than the ground truth resolution of 0.025 cycles per voxel.

### 3.2 Refinement against the known structure at the correct resolution in each iteration yields reconstructions without over-fitting

In order to confirm that the observation of over-fitting is due to the accumulation of noise, we performed the same refinement using the ground-truth (the known structure) as the reference volume for each refinement iteration. Identical synthetic data and refinement procedures were used, however the reconstructed half maps were replaced with the synthetic template structure filtered to the current resolution at each iteration. Little-to-no over-fitting was observed beyond the expected cut-off in each segment, both as measured by FSC extension and based on visible features (Fig. 2, idealised reconstruction). Note that all of the FSC curves in this case fall off steeply to near zero, indicating minimal residual correlation at higher resolutions.

### 3.3 Over-fitting is substantially mitigated by application of the SIDESPLITTER algorithm, restricting the reconstruction to regions that can be assigned with confidence

After confirming that we could reproduce the over-fitting issue under controlled conditions with a known ground-truth, we attempted to mitigate against it using the SIDESPLITTER algorithm. We used identical synthetic data and an identical refinement process, however the SIDESPLITTER filter was applied to the half maps between iterations, and the corresponding output used for the next iteration of the refinement. The results were comparable to the use of the ground-truth from the point of view of visible features, in that little over-fitting was observed beyond the expected cut-off in each segment. The FSC curves revealed greater retention of noise at higher resolutions than for the ground-truth case, but substantially less than was the case for the original refinement (Fig. 2, SIDESPLITTER reconstruction. Compare in particular the orange and yellow curves between the SIDESPLITTER and gold-standard refinements.) This implies that SIDESPLITTER mitigated but did not completely alleviate over-fitting for synthetic data.

### 3.4 Application of the SIDESPLITTER algorithm successfully mitigates against over-fitting in an experimental dataset with a low local resolution detergent micelle

Having confirmed the benefit of the approach in principle, we then set out to confirm that it was applicable in practice to experimental data. Two experimental conditions were considered, that in which the region of lower local resolution in question is known to lack consistent structure between particles, and that in which there is known and quantifiable heterogeneity between subpopulations of particles. In the first case, we tested SIDESPLITTER using the recent structure of human amino acid transporter LAT1 bound to CD98 and an antibody fragment within a detergent micelle (EMD-9849; Lee *et al*, 2019). Within the micelle the individual detergent molecules are expected to adopt unrelated positions away from the immediate environment of the protein, however in the original reconstruction, some structure remains apparent within the micelle which is presumably due to over-fitting of noise. In the original work, subtraction of the micellar region resulted in a higher-resolution structure of the protein. The application of SIDESPLITTER refinement to the original data resulted in a structure with a substantial reduction in both the power of, and the features within, the micelle (Fig. 3A, S3), and an improvement in resolution of the protein that was comparable to the micelle-subtraction approach.

**Figure 3:**
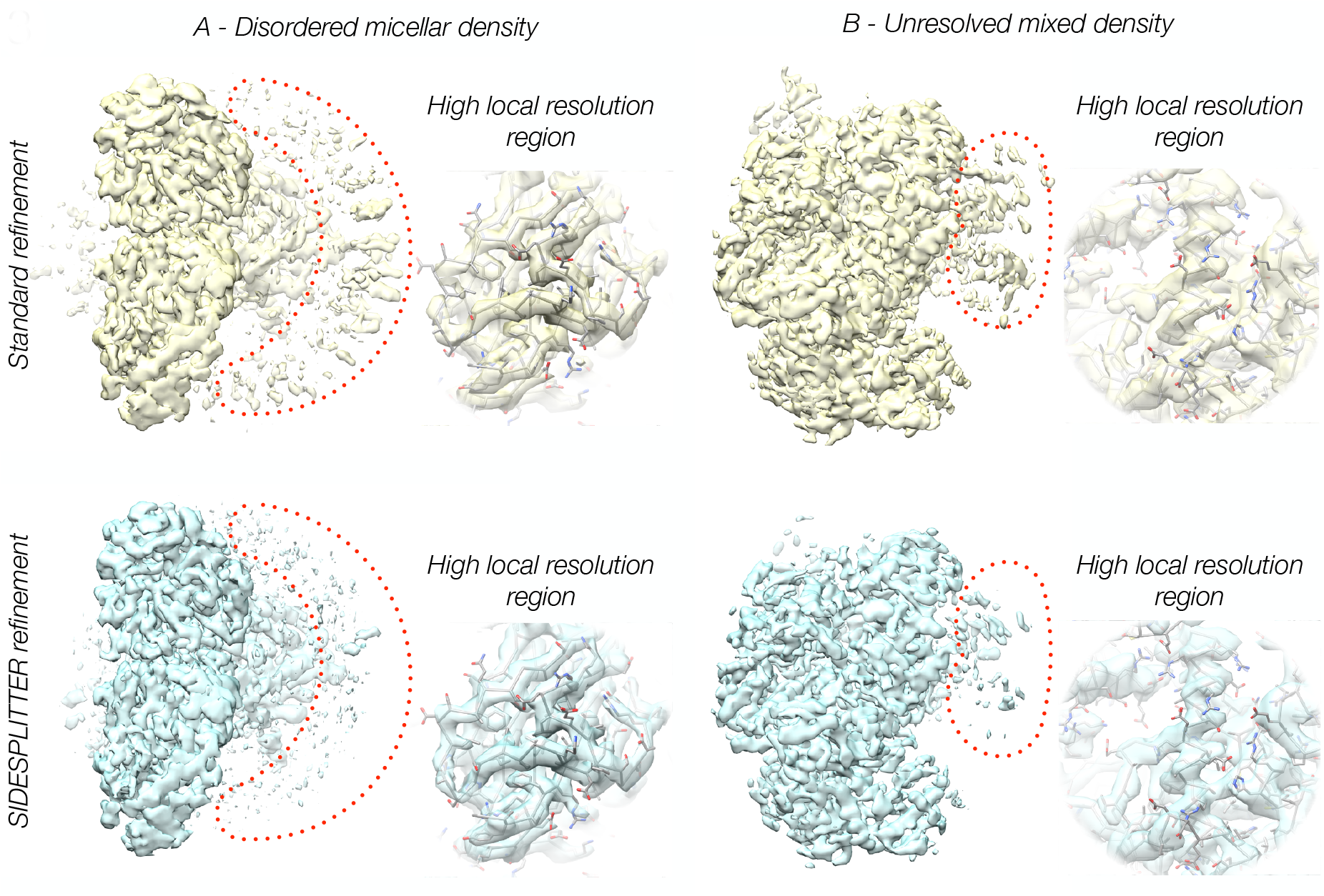
Refinement of experimental datasets with substantial regions of known low local resolution / SNR shows that SIDESPLITTER refinement suppresses features within regions of low local resolution, and improves the quality of the final density. Comparison between the results of standard and SIDESPLITTER refinement for (A) EMD-9849, and (B) EMD-4038. Volumes are shown as transparent surfaces. Regions of high local resolution are inset in each case, with the corresponding PDB (PDB-ID 6JMQ and PDB-ID 5LD2 respectively) fitted into the densities in question, demonstrating that signal is retained to high resolution in regions with a high local signal to noise ratio.

### 3.5 Application of the SIDESPLITTER algorithm successfully down-weights regions of an experimental dataset known to correspond to multiple conformational states

In a second experimental case, a RecBCD dataset (EMD-4038; Wilkinson *et al*, 2016), we ran a single reconstruction combining particles that had previously been split into four classes, in which movement of one domain is evident. We would expect a loss of features within these regions if our approach is successful. The application of SIDESPLITTER refinement to this data resulted in a reduction of visible features and power within this region exactly as expected (Fig. 3B).

### 3.6 The SIDESPLITTER algorithm does not degrade the final resolution limit attained, and will yield higher resolution in cases in which over-fitting is severe

For both experimental applications, and in other tests performed to date (data not shown), density from SIDESPLITTER refinement appears to be clearer and cleaner than that from standard refinement (Fig. 3). For the dataset exhibiting unresolved heterogeneity, the apparent resolution by FSC was identical to that in the case of standard refinement, implying that SIDESPLITTER is not derogatory to the overall resolution attained. For the micellar case, the resolution according to FSC 0.143 is higher than in the case of standard refinement (3.62 Å versus 3.76 Å, respectively), and equivalent to that from a dataset in which subtraction of the micelle has been performed (Lee *et al*, 2019).

## 4. Discussion

Over-fitting within macromolecular structures is particularly pernicious, as it undermines the interpretation of biological function and activity. If, within regions of a reconstruction, the noise dominates, it may be mistaken for signal, rendering any interpretation necessarily flawed. The twin problems of the resolution at which a reconstruction remains interpretable, and of variable local “resolution” or SNR, have been heavily investigated within the single particle analysis field. Independent half-set refinement, in which over-fitting is mitigated against during the refinement process (Grigorieff, 2000; Scheres & Chen, 2012; Henderson *et al*, 2012), local resolution measurement (Cardone *et al*, 2013, Kucukelbir *et al*, 2014) and local resolution filtering (Cardone *et al*, 2013; Vilas *et al*, 2018a), pursued after the refinement process, have been widely adopted to avoid over-interpretation of reconstructed densities.

Here we have shown that, despite these advances, over-fitting during the refinement process within areas of low local resolution / SNR remains problematic during independent half-set refinement. The two half sets contributing to the reconstructions can be kept independent; however, the noise within the resulting reconstructions will not be uncorrelated. Despite the separation of the two sets of particle images, certain characteristics are shared between the sets, including the regions of lower local resolution (and corresponding local high-resolution noise), the orientation distribution, initial model, and the mask used in refinement. This means that density corresponding to noise will tend to accumulate similarly, even if independently, on each side of a split refinement. This process leads inexorably to over-fitting in more poorly-resolved regions of the reconstructed density through the positive feedback process of iterative refinement, noise being aligned against noise. Such over-fitting cannot be entirely mitigated against after the refinement, as the incorporated noise becomes indistinguishable from signal. Affected regions will have higher apparent SNR, and exhibit higher apparent resolution, than should be the case given the underlying data, and flawed interpretation of such structures is a very real risk.

Remedies for this using some form of local filtering have been proposed previously, and basic implementations provided (Grigorieff, 2008) for simple versions of weighting approaches with user intervention. In looking for an algorithm to allow automatic and unbiased weighting along these lines, the major (and non-trivial) problem is to maintain the independent nature of the split refinement. The application of windowed local-resolution filters cannot maintain this independence, as the shared signal within local windows must necessarily become correlated, and therefore such filters cannot be compatible with an independent split refinement (Supp. Fig. 1). We have overcome this issue by creating a local SNR filter suitable for independent refinement of a split dataset. SIDESPLITTER, based on a modified local SNR filter that minimises the residual noise within the two reconstructions (Ramlaul *et al*, 2019), maintains the independence of the two sides of a split refinement during the refinement process by taking account of only the global noise distribution between them.

We have shown that SIDESPLITTER effectively mitigates over-fitting both in synthetic situations, where we have explicitly generated and measured over-fitting, and in experimental data with known over-fitting problems which have previously been mitigated by the manual interventions of particle sorting and density subtraction. SIDESPLITTER refinement has shown demonstrable improvements in the output density in situations with large regions of lower local resolution, and the reduction of over-fitting has been shown to increase the overall resolution in particularly egregious cases, where regions of low local resolution make up a substantial portion of the refined density. We believe that the SIDESPLITTER approach will be of benefit to the field during any refinement in which there is a notable variation in local resolution within the volume.

## Abbreviations

2D/3D: 2/3-Dimensional
Cryo-EM: Electron Cryo-Microscopy
EM: Electron microscopy
FSC: Fourier shell correlation
LAFTER: Local Agreement Filter for Transmission EM Reconstructions
SNR: Signal to noise ratio

## Acknowledgments

The authors would like to thank; R Ayala, K Chen, EYD Chua, Y Lee, M Wilkinson, O Willhoft, DB Wigley and X Zhang for providing data for beta testing of SIDESPLITTER on problematic electron microscopy datasets. We also thank T Nakane and SHW Scheres for providing the RELION 3.1 alpha version for refinement testing.

---

**Supplementary Figure 1:**
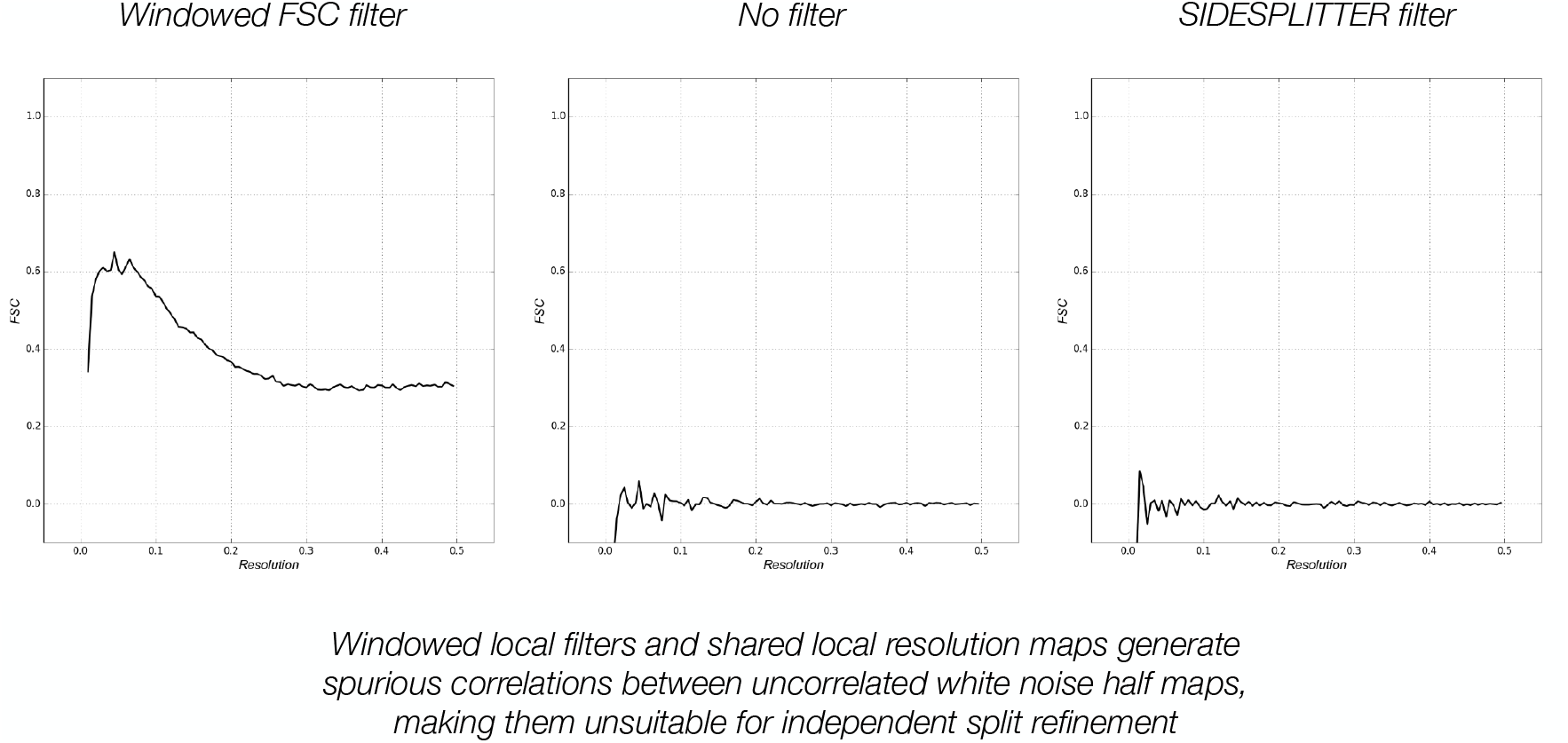
Windowed local filters and shared local resolution maps are unsuitable for independent split refinement as they generate spurious correlations. Half-set FSC curves after the application of a windowed FSC filter (BLOCRES/BLOCFILT) (left), without any processing (centre), and after the application of SIDESPLITTER (right) to the same uncorrelated white noise densities. The curves oscillate around zero where a correlation has not been generated by the local filtering process. The BLOCRES/BLOCFILT filter introduces correlation over the entire resolution range. By contrast, the SIDESPLITTER filter introduces no discernible correlation.

**Supplementary Figure 2:**
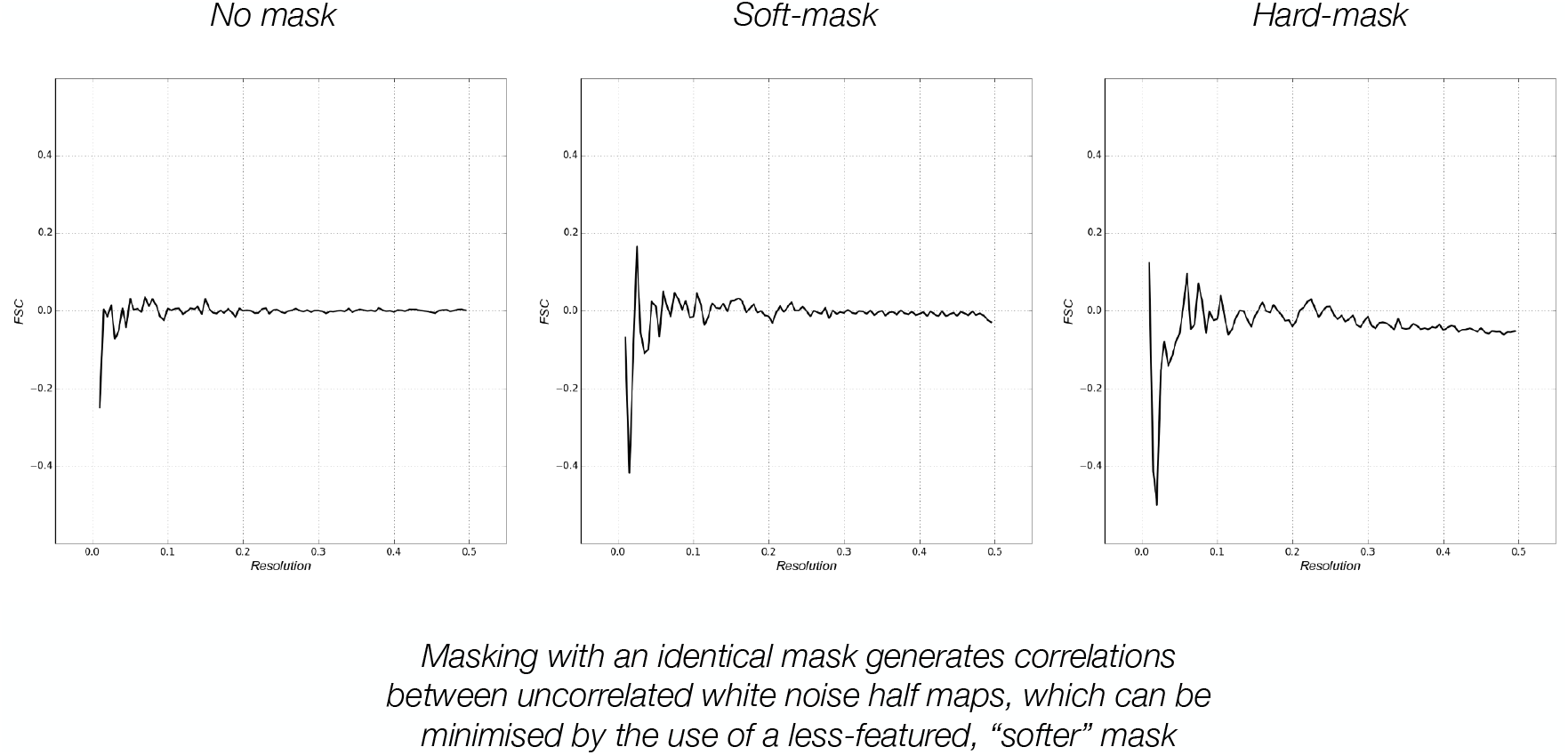
Shared solvent masks also generate spurious correlations, although the effect is not pronounced, especially for soft masks. Resulting half-set FSC curves after the application of no mask (left), a soft mask (centre), and a relatively sharp mask (right) to the same uncorrelated noise densities. The curves oscillate around zero where a correlation has not been generated by the local filtering process. In the unmasked noise half-maps, oscillations indicating correlations are much smaller in magnitude compared to the masked half-maps. The hard-masked maps in particular display aggravated correlations at low resolution and some correlation at high resolution. The correlations in the soft-masked maps are smaller and occur only at low resolution, but they are still noticeable, indicating that artefactual correlations due to masking can be minimised, but not abrogated, by using softer masks.

**Supplementary Figure 3:**
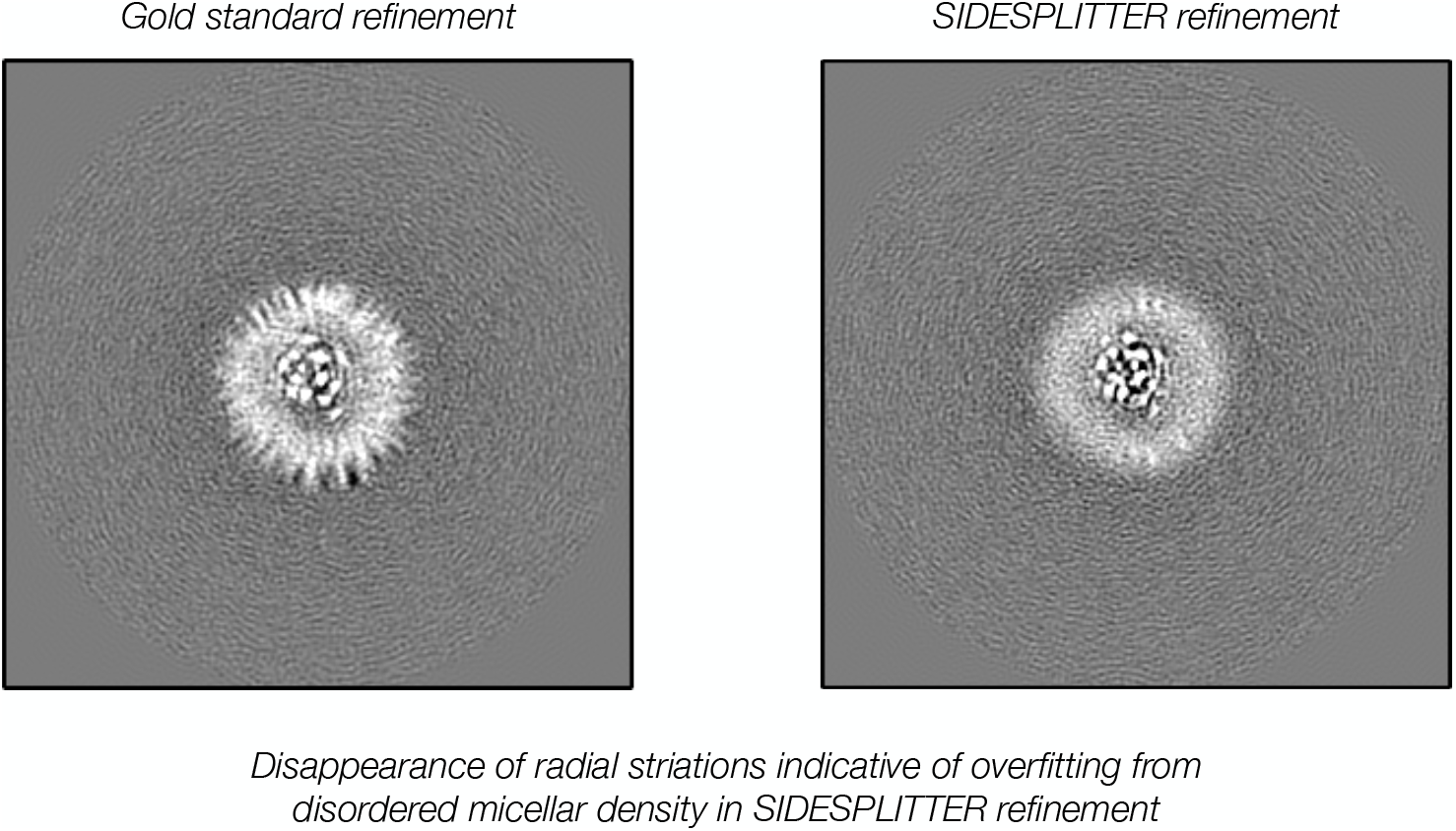
SIDESPLITTER refinement results in the supression of striations due to overfitting. Sections through the micellar density showing visible evidence of overfitting in standard refinement, which are supressed on SIDESPLITTER refinement.

